# Paroxysmal Slow Wave Events as a diagnostic and predictive biomarker for post-traumatic epilepsy

**DOI:** 10.1101/2024.10.14.618064

**Authors:** Gerben van Hameren, Pooyan Moradi, Hamza Imtiaz, Ellen Parker, Saara Mansoor, Laith Al Hadeed, Mohammed Albitar, Ziad Alhosainy, Alon Friedman

## Abstract

Traumatic brain injury (TBI) is a major global health concern, affecting more than 40 million people annually. While most cases are mild and present with light symptoms, repeated mild injuries can result in delayed brain pathologies, including cognitive decline, neuropsychiatric complications, and post-traumatic epilepsy (PTE). PTE refers to recurring, unprovoked seizures occurring at least one week after TBI. While the link between moderate to severe TBI and PTE is well established, the epileptogenesis after repetitive mild TBI (rmTBI) is seldom studied.

Currently, there are no biomarkers to identify those at risk of developing PTE, and its diagnosis is challenging. Here, we used a rat model to study PTE following rmTBI and assessed human EEG data to identify potential biomarkers for PTE.

We employed a closed head TBI model to induce rmTBI, and recorded brain activity using electrocorticography (ECoG) between 2- and 6-months post-injury. Behavioral assessments and post-mortem analysis were also conducted. In humans, we analyzed EEG recordings from the Temple University database to investigate the potential of EEG-derived features for diagnosing PTE.

At 6 months post injury, 70% of rmTBI animals developed PTE, compared to 22% in the control group (P=0.01). While neurological assessments following injury did not predict PTE, paroxysmal slow wave events (PSWEs) were found to be a reliable biomarker for PTE prediction. In humans, the percentage time in PSWEs was significantly elevated in PTE patients with epileptiform activity.

In conclusion, we suggest PSWEs as a non-invasive, cost-effective biomarker for PTE in rodents and human patients.

## Introduction

Traumatic brain injury (TBI) is a major global health concern, affecting more than 40 million people annually *(1)*. Falls, car accidents, and sport-related injuries are among the most common causes of TBIs. Cumulative data shows that while the incidence of TBI increases over the years, the number of TBI-related deaths has decreased *(2)*. This translates to a higher rate of patients who survive, but are at risk of long-term complications, including physical, behavioral, emotional, neurological, and cognitive disabilities *(3, 4)*.

Mild TBIs are the most common type of TBI and a leading cause of neurological disorders globally. Particularly, repetitive mild TBI (rmTBI) has been reported to elevate the risk for neuropsychiatric disabilities later in life. rmTBI is particularly common among athletes in high-impact sports (e.g., football, hockey, rugby, mixed martial arts) as well as among military personnel *(5–8)*. Additionally, the steady increased risk of falls among the aging population contribute to a greater incidence of rmTBI *(9)*.

Post-traumatic epilepsy (PTE) is a long-term TBI complication in which people experience seizures after the initial brain injury *(10)*. PTE is defined by unprovoked seizures occurring more than a week after brain injury. The onset of PTE varies between patients, but they typically suffer their first seizure between 1 and 2 years after the injury *(11–15)*. Little is known about the prevalence of PTE following mild TBIs, partly due to underreporting and poor follow-up, but it is likely lower compared to moderate-severe TBIs. However, because the prevalence of mild TBI cases is high andrepeated injury may increase the risk for PTE *(16)*, rmTBI still contributes significantly to the number of PTE patients.

Living with epilepsy drastically decreases one’s quality of life and is associated with comorbidities such as depression and cognitive decline *(17, 18)*. PTE can be difficult to diagnose, especially when seizures’ phenotype is atypical *(20)*. In addition, PTE can be drug-resistant, making it difficult to treat *(21)*. Despite advancements in understanding PTE pathophysiology, there are currently no reliable biomarkers for identifying patients at high risk of developing PTE. Predictive and diagnostic biomarkers are essential for the design and success of clinical trials, patient stratification, and the timely and quantitative assessment of target engagement and treatment efficacy *(22)*.

Quantitative brain activity data derived from electroencephalograms (EEGs) offer promise as a low-cost, non-invasive approach to predict PTE *(23)*. In previous work in patients with epilepsy, as well as in animal models of induced epilepsy, a pattern of intermittent, paroxysmal slowing of cortical activity was identified and termed paroxysmal slow wave events (PSWEs) *(24)*. Additionally, PSWEs were found more frequently 48 hours after the first seizure in patients who later developed epilepsy *(25)*, suggesting PSWEs as a potential biomarker of epileptogenesis.

In the current study, we aimed to characterize PTE in an rmTBI model in rats. In search of predictive biomarkers, we assessed neurological and cognitive changes in response to rmTBI and used longitudinal electrocorticography (ECoG) recordings to measure neural network changes, PSWEs and detect epileptiform activity. Lastly, we used the Temple University EEG dataset to analyze brain activity acquired from patients who developed epilepsy following TBIs and investigated the potential of PSWEs as a diagnostic biomarker for PTE.

## Results

### repetitive mild TBIs are linked to locomotion and cognitive impairments

rmTBI was induced in a total of 80 animals (for experimental timeline see Fig. 1A and Methods). rmTBI resulted in neurological changes consistent with previous work in our lab *(26)*. The earliest response to head injury were convulsive movements, most often in lower limbs and tail, which were observed in 29 out of 52 rats (Fig. 1B) and never in anaesthetized controls. Righting latency was significantly longer in rmTBI animals compared to controls (Fig. 1C). Following 4 impacts, rmTBI animals lost an average of 2.34% ± 4.23% of their body weight (8.62 g ± 15.62 g) compared to their baseline. In contrast, control animals gained an average of 4.39% ± 2.63% (16.32 g ± 9.82 g) over the same period (Fig. 1D). Three rmTBI rats were excluded from the study after severe weight loss following injury. Neurobehavioral assessment showed a progressive reduction in neurological severity score (NSS, see methods) in rmTBI animals compared to controls (Fig. 1E). At 3 weeks post injury, we assessed cognitive performance using Novel Object Recognition (NOR) test and Elevated Plus Maze (EPM). rmTBI animals did not vary significantly from controls in NOR test at 3 weeks post injuries (Fig. 1F). In contrast, anxiety index was significantly higher in rmTBI animals compared to sham controls (0.86 ± 0.09 vs 0.79 ± 0.08, in rmTBI vs control animals respectively; Fig. 1G), suggesting higher anxiety.

**Figure 1.**
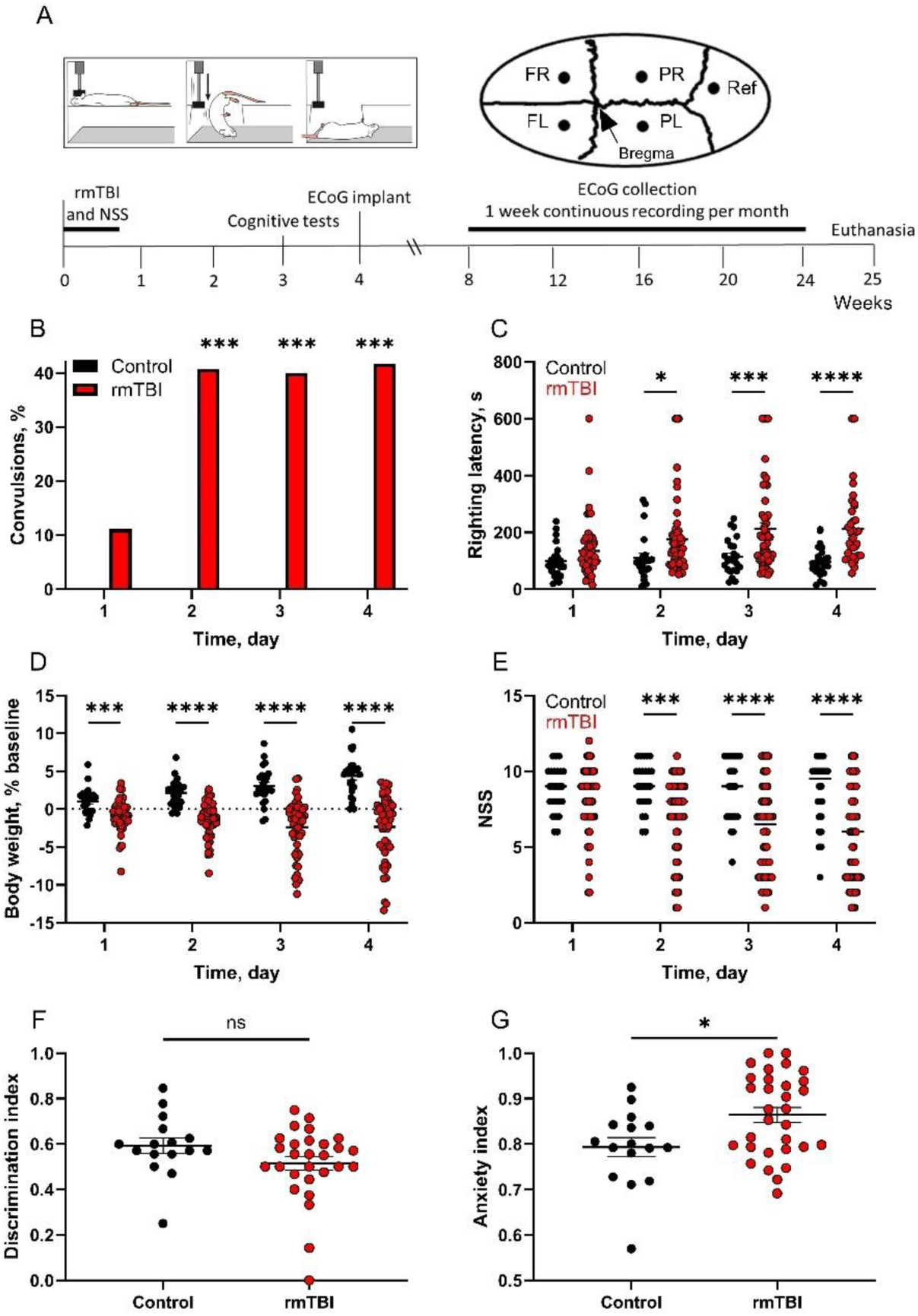
Neurobehavioral outcomes following rmTBI. **A)** Experimental timeline: Animals underwent repetitive mild TBIs, followed by ECoG electrode implantation 4 weeks post-injury. ECoG recordings were continuously collected from 2 to 6 months after electrode implantation. **B)** rmTBI animals show convulsions acutely after head injury, whereas controls do not (1^st^ hit, P=0.08, control n=25 vs rmTBI n=55; 2^nd^ hit, P=0.0002, control n=25 vs rmTBI n=55; 3^rd^ hit, P=0.002, control n=25 vs rmTBI n=50; 4^th^ hit, P=0.002, control n=25 vs rmTBI n=36; Pearson’s chi-square tests). **C)** Righting latencies are longer after rmTBI, compared to controls (1^st^ hit, P=0.18, control n=25 vs rmTBI n=55; 2^nd^ hit, P=0.03, control n=25 vs rmTBI n=55; 3^rd^ hit, P=0.0009, control n=25 vs rmTBI n=50; 4^th^ hit, P<0.0001, control n=25 vs rmTBI n=36; Sidak’s multiple comparisons tests. **D)** Rats lose weight after rmTBI, while control animals gain weight (1^st^ hit, P=0.0004, control n=25 vs rmTBI n=55; 2^nd^ hit, P<0.0001, control n=25 vs rmTBI n=54; 3^rd^ hit, P<0.0001, control n=25 vs rmTBI n=54; 4^th^ hit, P<0.0001, control n=25 vs rmTBI n=53, Sidak’s multiple comparisons tests). **E)** NSS was significantly lower in rmTBI animals after 2^nd^ hit compared to controls (1^st^ hit, P=0.38, control n=25 vs rmTBI n=55; 2^nd^ hit, P=0.0002, control n=25 vs rmTBI n=54; 3^rd^ hit, P<0.0001, control n=25 vs rmTBI n=54; 4^th^ hit, P<0.0001, control n=25 vs rmTBI n=53, Sidak’s multiple comparisons tests). **F)** Performance in NOR test (discrimination index: visits to novel object) is not different between rmTBI and control animals at 3 weeks post-injury (P=0.1, control n=16, rmTBI n=28; Mann-Whitney test). **G)** Anxiety index was significantly higher in rmTBI animals compared to sham controls at 3 weeks post-injury (P=0.03, control n=16, rmTBI n=31, Mann-Whitney test).

### Seizure-like events increase in frequency over time and primarily recorded in frontal electrodes

ECoG recordings of cortical activity from 4 epidural electrodes, 2 - 6 months after injury in rmTBI-exposed rats were semi-automatically analyzed for seizure-like events (SLEs; Fig. 2A). SLE frequency was higher in rmTBI animals compared with controls (38.76 ± 8.06 vs 11 ± 5.05 SLEs/day in rmTBI and control animals, respectively, P=0.01; Fig. 2B). SLEs’ characterization demonstrated that SLEs are significantly longer in duration in rmTBI animals (19.58 ± 0.3 vs.15.55 ± 0.63 in rmTBI animals and controls respectively, P<0.0001; Fig. 2C). Notably, a duration threshold of >20 seconds identified as the optimal discriminator between SLEs in rmTBI and control animals (P<0.0001; Pearson’s chi-square test). Additionally, SLEs in rmTBI animals were more likely to be recorded in two or more electrodes simultaneously (22.4 ± 12.3% vs 7.8 ± 8.74% in rmTBI animals and controls respectively, P=0.0001; Fig. 2D).

**Figure 2.**
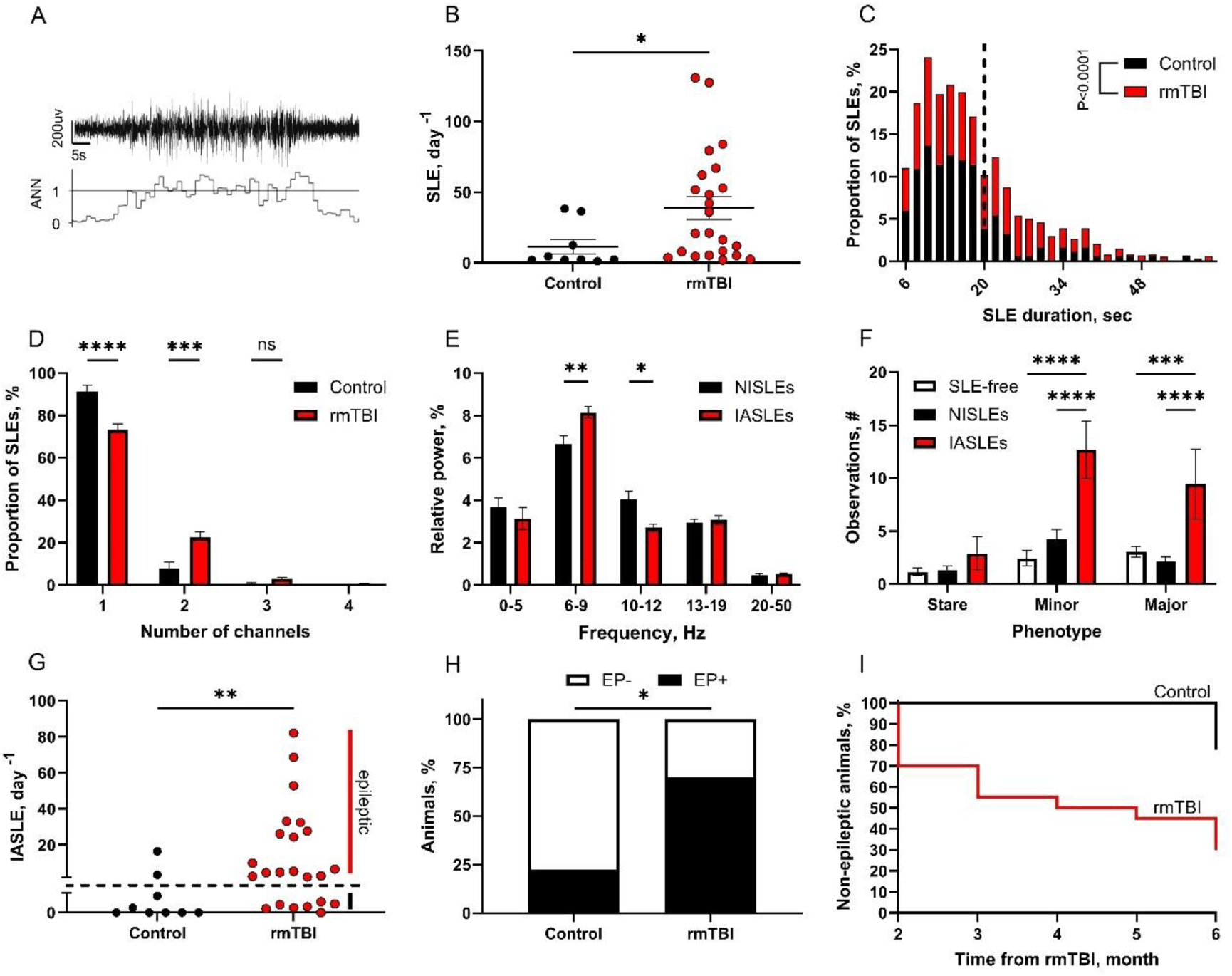
rmTBI is linked with a higher occurrence of SLEs. **A)** Example of ECoG recording during a “seizure-like-event” (SLE, upper trace). NVision seizure detection algorithm shows an artificial neural network (ANN) score above 1 during the SLE (lower trace). **B)** The SLE frequency in rmTBI animals was higher compared to controls (P=0.01; control n=9 vs rmTBI animals n=23; Mann-Whitney test). **C)** In rmTBI animals, the mean duration of SLEs was significantly greater than in controls (P<0.0001; Mann-Whitney test). In addition, relative frequency analysis demonstrated that rmTBI animals had a higher incidence of prolonged (>20s) SLEs than controls. **D)** In rmTBI animals, a significantly larger proportion of SLEs were simultaneously detected by two electrodes compared to controls (P<0.0001; Two-way ANOVA). Conversely, the occurrence of SLEs detected by only one electrode was notably higher in controls than in rmTBI animals (P<0.0001; Two-way ANOVA). **E)** IALSLEs showed significantly higher mean relative power in 6-9 Hz and lower power in 10-12 Hz frequencies (6-9 Hz: P=0.004, 10-12 Hz: P=0.01; Two-way ANOVA). **F)** The mean number of motor behavioral abnormalities (minor and major motor movements) was notably higher in IASLEs compared to observations from SLE-free and NASLEs zones (Stare: P=0.98 in NISLEs vs SLE-free, P=0.54 in IASLEs vs SLE-free, P=0.57 in IASLEs vs NISLEs; Minor movements: P=0.29 in NISLEs vs SLE-free, P<0.0001 in IASLEs vs SLE-free, P<0.0001 in IASLEs vs NISLEs; Major movements: P=0.74 in NISLEs vs SLE-free, P=0.0005 in IASLEs vs SLE-free, P<0.0001 in IASLEs vs NISLEs; SLE-free segments n=19, NISLEs n=29, IASLEs n=9, collected from 8 animals; Two-way ANOVA). **G)** Peak occurrence per day of IASLEs was higher in rmTBI animals compared to controls (P=0.003, control n=9 vs rmTBI n=23; Mann-Whitney test). **H)** At six months, the percentage of animals classified as epileptic (EP+) was significantly higher in the rmTBI group (control n=9, rmTBI n=20, P=0.01; Pearson’s chi-square test). **I)** Percentage of non-epileptic animals showed a progressive decline overtime in rmTBI group (2 months: 0% vs 34.78%, control n=9, rmTBI n=23; 3 months: 0% vs 47.82%, control n=9, rmTBI n=21; 4 months: 0% vs 52.17%; 5 months: 0% vs 56.52%, control n=9, rmTBI n=20; 6 months: 22.22% vs 69.56%, control n=9, rmTBI n=20).

Based on these observations, we identified prolonged, multifocal “injury-associated SLEs” (IASLEs; ≥20s in duration and recorded in more than 1 electrode) and “non-injury SLEs (NISLEs). To assess signal characteristics, we performed spectral analysis and found that IASLEs exhibit increased relative power in lower frequency bands compared to NISLEs. Specifically, IASLEs showed significantly higher relative power in the 6-9 Hz range (8.14 ± 1.69 vs. 6.64 ± 1.87 in IASLEs and NISLEs respectively; P=0.004; Fig. 2E). Conversely, NISLEs demonstrated greater relative power in the 10-12 Hz frequency bands (4.07 ± 1.67 vs. 2.71 ± 1.05 in NISLEs and IASLEs respectively, P=0.01). The number of observed behavioral abnormalities during IASLEs was also significantly higher compared to those occurring during randomly selected SLE-free zones (see methods) as well as in NISLEs (Minor abnormal movements: 12.66 ± 2.69 in IASLEs vs 2.44 ± 0.71 in SLE-free and 4.27 ± 0.87 in NISLEs; Major abnormal movements: 9.44 ± 3.13 in IASLEs vs 3.05 ± 0.52 in SLE-free and 2.17 ± 0.41 in NISLEs; Fig. 2F).

Based on our observations, we propose IASLEs (and not NISLEs) as potential indicative feature of pathological epileptiform activity, more likely to develop in the rmTBI-exposed rat. The peak occurrence of IASLEs was indeed significantly higher in rmTBI animals than in controls (Fig. 2G) and a frequency of two or more IASLEs per day appeared to effectively distinguish rmTBI and control animals (Fig. 2G, P<0.01, Pearson’s chi-square test). In this model, 70% of rmTBI animals were identified as epileptic at 6 months post injuries, compared to 22% of sham controls (Fig. 2H). Retrospectively, a progressive increase in the percentage of epileptic animals was observed in rmTBI group compared to the sham controls (Fig. 2I). Notably, two (out of nine) control animals developed epilepsy at 6 months after repeated exposure to isoflurane.

To further characterize PTE, we analyzed IASLEs’ localization, daily variation and progression with time. A majority of IASLEs were detected in frontal electrodes (80 and 85% in left and right frontal respectively; Fig. 3A), whereas IASLEs recorded by the parietal electrodes were less common (12 and 22% in left and right parietal respectively). Note that the same IASLEs were often detected by two electrodes simultaneously. Some IASLEs (17%) were first recorded in the frontal cortex and seems to propagate to the posterior electrode with a velocity of 0.67 ± 1mm/s. IASLEs first recorded in the parietal electrodes and apparently propagated to the frontal electrodes were less common (5%; Fig. 3A). Mean IASLE frequency during the active (lights-off) period was higher compared to resting (lights-on) period (1.48 ± 0.18 vs 0.92 ± 0.52; Fig. 3B). Interestingly, the mean duration and frequency of IASLEs increased over time (Fig. 3C, D). Notably, a subset of epileptic rmTBI animals (8 out of 14) showed increased frequency of IASLEs between 2-6 months and were therefore referred as “progressive PTE” (Fig. E, F). In contrast, the remaining 6 animals showed either stable or decreased IASLE occurrence, categorized as "non-progressive PTE".

**Figure 3.**
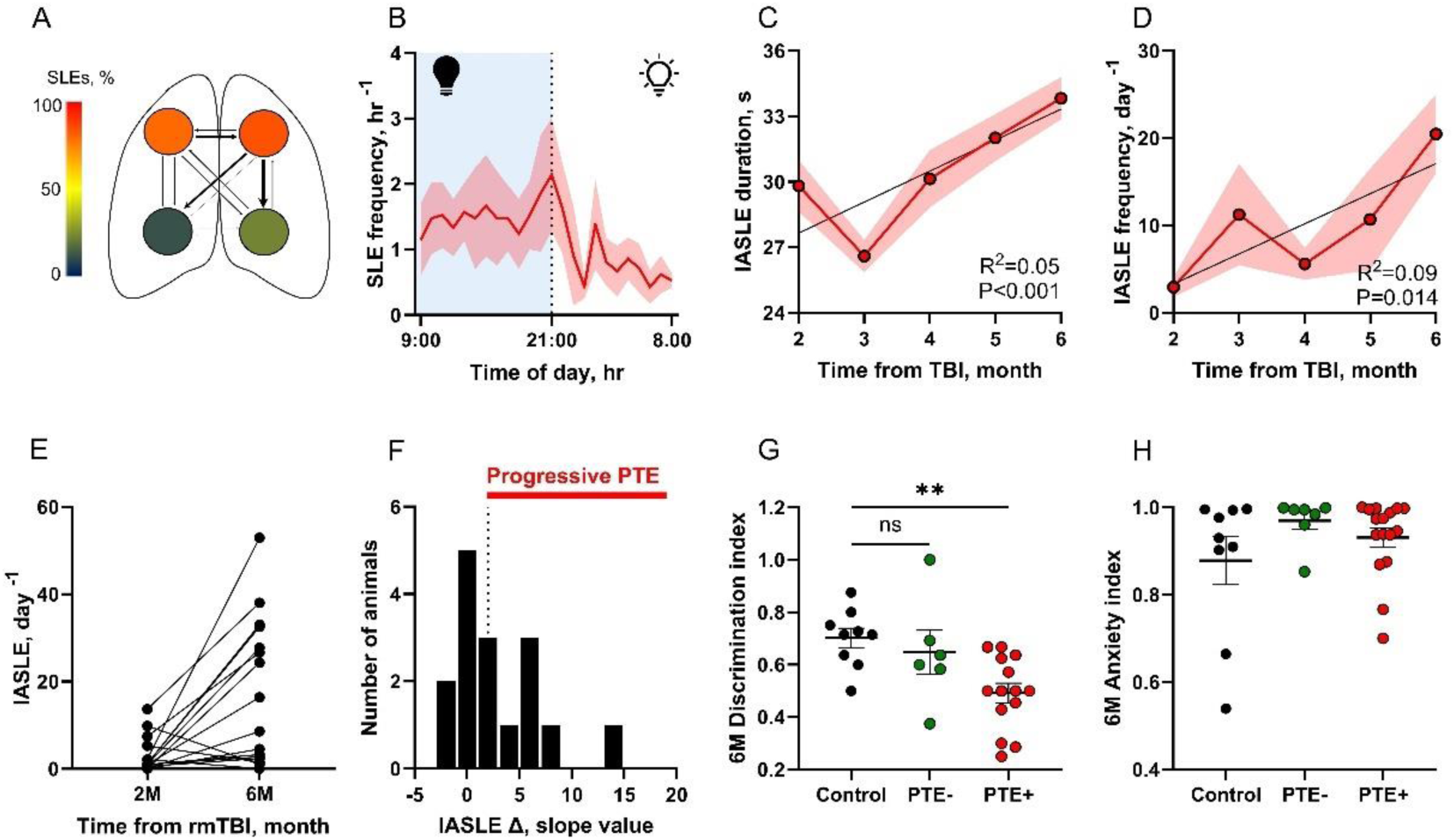
Characterisation of PTE following rmTBI. **A)** Spatial analysis of IASLE detection (n=861) by the four ECoG recording electrodes. IASLEs with a detectible propagation pattern are indicated by arrows, with the arrow size indicating their relative frequency. **B)** The average number of IASLEs per hour during the active (lights-off) period was significantly greater than during the resting (lights-on) period (n=21, P=0.005, Mann-Whitney test) **C)** IASLEs’ duration increased over time in epileptic animals (R^2^=0.05, P<0.001; simple linear regression). **D)** IASLEs’ occurrence per day also increased over time (R^2^=0.09, P=0.014; simple linear regression). **E)** IASLEs’ occurrence per day at 2 vs 6 months post rmTBI. Change in IASLE’s occurrence was heterogeneous in epileptic animals. **F)** Distribution of animals with progressive PTE (slope >1, n=8) and non-progressive PTE (slope <1, n=6; simple linear regression). **G)** Cognitive testing was performed on a subset of animals at 6 months post rmTBI. PTE+ but not PTE- animals, performed worse in NOR test compared to sham control (control n=9 vs PTE- n=6, P=0.55 and vs PTE+ n=14, P=0.002; Kruskal-Wallis test followed by Dunn’s post-hoc test). **H)** No significant difference was found in EPM anxiety index between groups (control n=9 vs PTE- n=6, P=0.22 and vs PTE+ n=14, P=0.80; Kruskal-Wallis test followed by Dunn’s post-hoc test).

At 6 months post rmTBI, animals that developed PTE (PTE+) exhibited significantly reduced performance in the NOR test compared to sham controls (0.49 ± 0.03 vs. 0.7 ± 0.03 in sham control vs rmTBI animals respectively, P=0.002; Fig. 3G). This suggests memory impairment as a co- morbidity associated with PTE. Differences in anxiety index were no longer measured at 6 months after injury (Fig. 3H).

### NSS and cognitive tests are not reliable predictors of PTE

We next asked whether neurological measures during the week of rmTBI can predict the development of PTE in the following months. No differences were found between animals classified with or without PTE in the presence or duration of acute, post-injury convulsive movements (Fig. 4A). Similarly, righting latency (Fig. 4B), weight loss (Fig. 4C), or NSS assessments (Fig. 4D) were no reliable predictors of PTE. While some epileptic animals exhibit memory decline as a PTE co-morbidity at 6 months, their NOR performance at 3 weeks does not reliably predict which animals will develop epilepsy later (Fig. 4E). EPM performance at 3 weeks after injury did not predict PTE development either (Fig. 4F). This lack of predictive power extends to discriminating PTE with progressive and non-progressive features (Fig. 4G-L).

**Figure 4.**
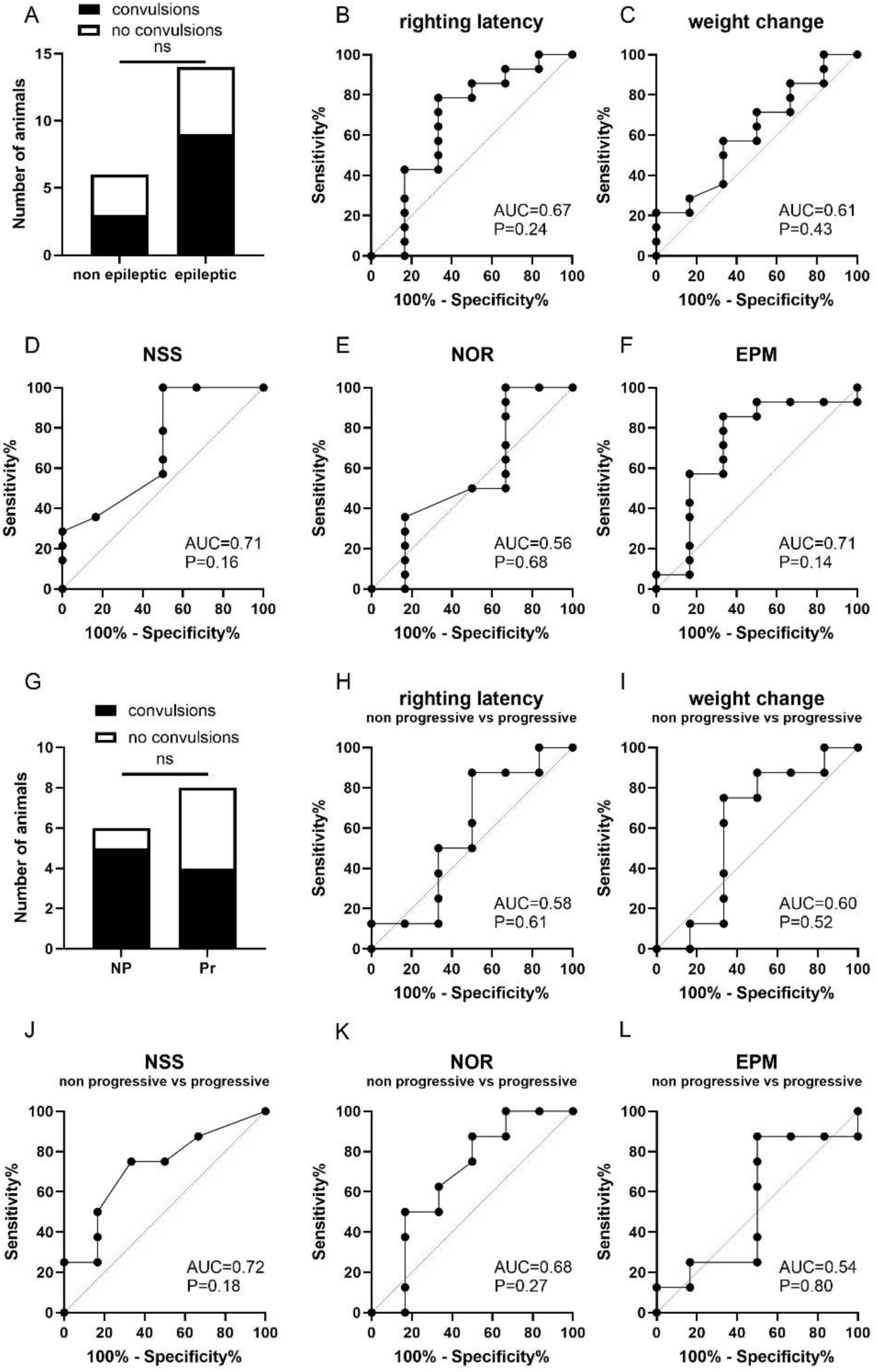
Neurobehavioral changes within 3 weeks post rmTBI do not predict PTE in animals. **A)** The percentage of animals with a history of post-injury convulsions did not differ between PTE- and PTE+ animals (P>0.67, Fisher’s exact test). **B)** Righting latency after the last hit does not predict PTE (P=0.24, ROC analysis). **C)** Weight change after the last hit does not predict PTE (P=0.43, ROC analysis). **D)** NSS scores after the last hit does not predict PTE (P=0.16, ROC analysis). **E)** NOR performance at 3 weeks post-rmTBI does not predict PTE (P=0.68, ROC analysis). **F)** EPM performance at 3 weeks post-rmTBI does not predict PTE (P=0.14; ROC analysis). **G)** The percentage of animals with post-injury convulsions did not differ between progressive and non-progressive PTE animals (P=0.37, Fisher’s exact test). **H)** Righting latency after the last hit does not predict progressive PTE (P=0.61, ROC analysis). **I)** Weight change after the last hit does not predict progressive PTE (P=0.52, ROC analysis). **J)** NSS after the last hit does not predict progressive PTE (P=0.18, ROC analysis). **K)** NOR performance at 3 weeks post-rmTBI does not predict progressive PTE (P=0.27, ROC analysis). **L)** EPM performance at 3 weeks post-rmTBI does not predict progressive PTE (P=0.8, ROC analysis). For PTE- vs PTE+ tests: control n=9, PTE- n=6, PTE+ n=14. For progressive vs non-progressive PTE tests: control n=9, non-progressive n=6, progressive n=8.

### PSWEs are more frequent in rmTBI animals with PTE

When exploring the features of the ECoG recordings, the power spectrum of the rmTBI rats demonstrated a higher reletive power between 1 and 5.5 Hz and increased area under the curve (AUC) in delta band compared with sham controls (Fig. 5A). It was previously shown that EEG/ECoG slowing in epileptic rodents and humans is due to the occurrence of short (>10 sec), paroxysmal events of cortical slowing (median power frequency <5 Hz), referred as PSWEs (Fig. 5B) *(24)*. Here, we assessed if PSWEs are also detectable following rmTBI in our recordings. PSWE occurrence peaked at the start of the rats’ inactive phase across all groups (Fig 5C). However, PTE+ animals showed increased PSWE occurrence compared to PTE- and sham controls at 2 months post rmTBI (8.22 ± 1.22, 3.71 ± 0.5 and 4.16 ± 0.65 in PTE+, PTE- and sham controls respectively). PSWEs were predominantly observed in ECoG electrodes implanted over the parietal cortex (36% in right and 39% in left parietal electrodes), while they were less frequently found in the frontal electrodes (12% in right and 13% in left frontal electrodes; Fig. 5D). Interestingly, the average PSWE occurrence at 6 months post rmTBI was significantly higher in PTE+ animals compared to sham controls (11.63 ± 3.27 vs 4.4 ± 0.68 in PTE+ and sham controls respectively; Fig. 5E).

**Figure 5.**
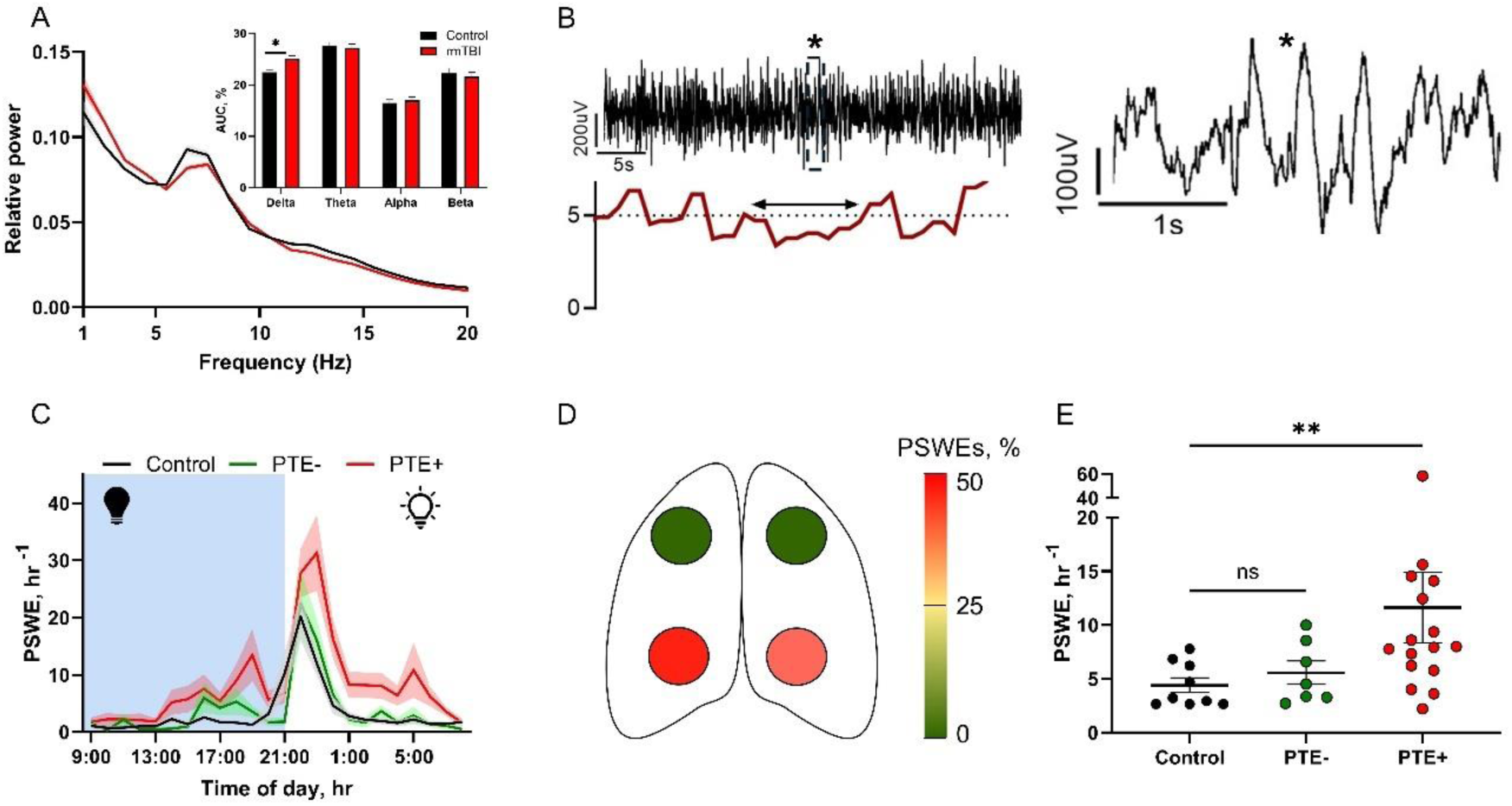
PTE is associated with increased number of PSWEs. **A)** Power spectrum of rmTBI and control rats. The inset displays the AUC for different frequency bands, with a significant increase in the delta band AUC in rmTBI rats (n=23) compared to controls (n=9; P=0.02, Mann-Whitney U test). **B)** Representative ECoG recording (top trace) showing a PSWE (double sided arrow), with the corresponding median power frequency (bottom trace) and a magnified 5-second segment (right). **C)** PSWE occurrence during the day shows a peak in the rat’s inactive phase (21.00-23.00). At 2 months post-rmTBI, PSWE occurrence is elevated in PTE+ animals during the day compared to both PTE- and sham controls (control n=9 vs PTE- n=7, P>0.9; control vs PTE+ n=16, P=0.04; PTE- vs PTE+, P=0.04; Kruskal-Wallis test followed by Dunn’s post-hoc test). **D)** Spatial analysis of PSWE detection by the four ECoG recording locations. **E)** Interestingly, average PSWE occurrence between 2-6 months post injury was significantly higher in PTE+ animals, but not in PTE- animals compared to controls (control n=9 vs PTE- n=7, P=0.64; control n=9 vs PTE+ n=16, P=0.008; Kruskal-Wallis test followed by Dunn’s post-hoc test).

### PSWEs are correlated with IASLEs and show predictive power for PTE

Next, we assessed the correlation between PSWEs and IASLE frequency at the same recording timepoint after rmTBI and found a significant association (Fig. 6A, p=0,019).. Since PSWEs were previously shown to predict epilepsy following a first seizure *(27)*, we then tested if PSWE frequency at our earliest timepoint (2 months after rmTBI) could predict which animals will develop PTE in later timepoints. PSWE frequency at 2 months showed significant predictive power for animals who developed PTE at 6 months post injuries (AUC=0.81, P=0.03; Fig. 6B) as well as for animals with progressive PTE (AUC=0.92, P=0.01; Fig. 6C). Interestingly, the occurrence of PSWEs at 2 months also correlated with memory impairment at 6 months post-rmTBI (R=-0.39, P=0.02; Fig. 6D).

**Figure 6.**
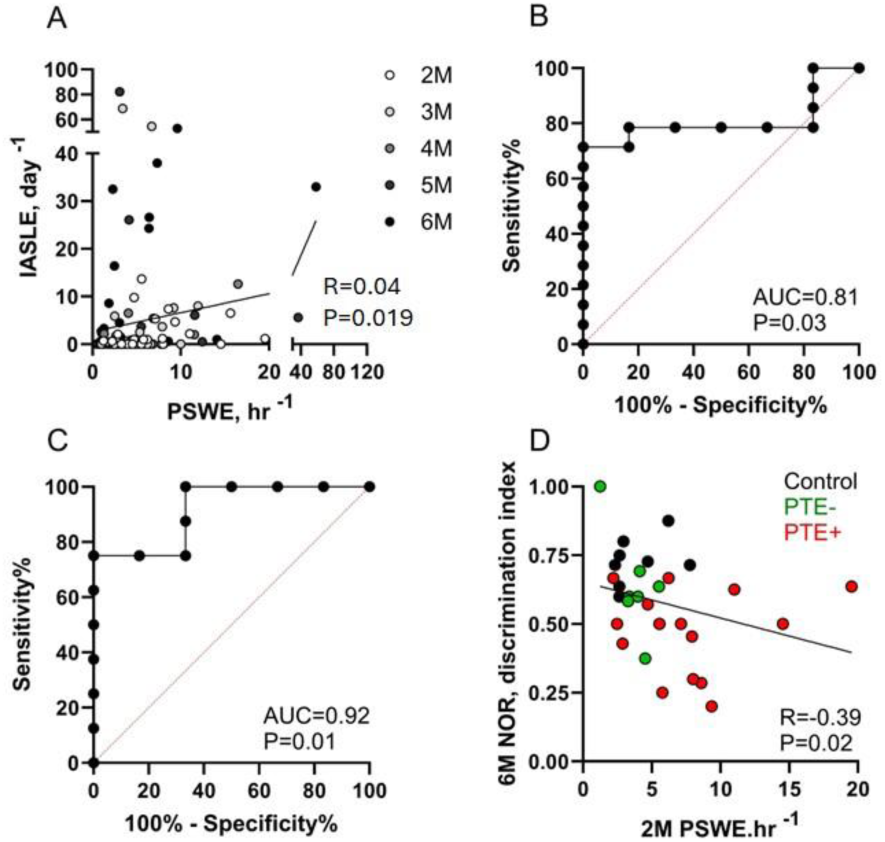
PSWEs are correlated with IASLE frequency and predict chance of PTE. **A)** PSWE frequency is correlated with IASLE frequency in the same ECoG recording (R=0.04, P=0.019; linear regression). **B)** PSWE occurrence per day at 2 months post rmTBIs show predictive power for animals that developed PTE later (PTE- n=6, PTE+ n=14, AUC=0.81, P=0.03; ROC analysis). **C)** PSWE occurrence per day at 2 months post rmTBIs also show predictive power for progressive PTE, notably with higher sensitivity (non-progressive PTE n=6, progressive PTE n=8, AUC=0.92, P=0.01; ROC analysis). **D)** PSWE frequency at 2 months was negatively correlated with NOR performance at 6 months post-rmTBI (control n=9, PTE- n=7, PTE+ n=15, P=0.02, R=-0.39; Pearson’s correlation test).

### rmTBIs induce region-specific neuroinflammation associated with PTE

To test if neuroinflammation could underlie epileptogenesis *(28)*, and if a seizure-onset zone could be detected, we analyzed markers of neuropathology in various brain regions. We assessed brains from a subset of sham controls and rmTBI-exposed animals, with and without PTE at the time of the perfusions (6 months post-injury). High variability in neuroinflammation markers was measured, especially in rmTBI animals. Both GFAP and IBA-1 were predominantly localized in the dentate gyrus. While GFAP expression remained unchanged (Fig. 7B), IBA-1 levels were significantly elevated in dentate gyrus region of TBI (both PTE- and PTE+) animals compared to sham controls (P=0.028 and P=0.022 respectively, Fig. 7C).

**Figure 7.**
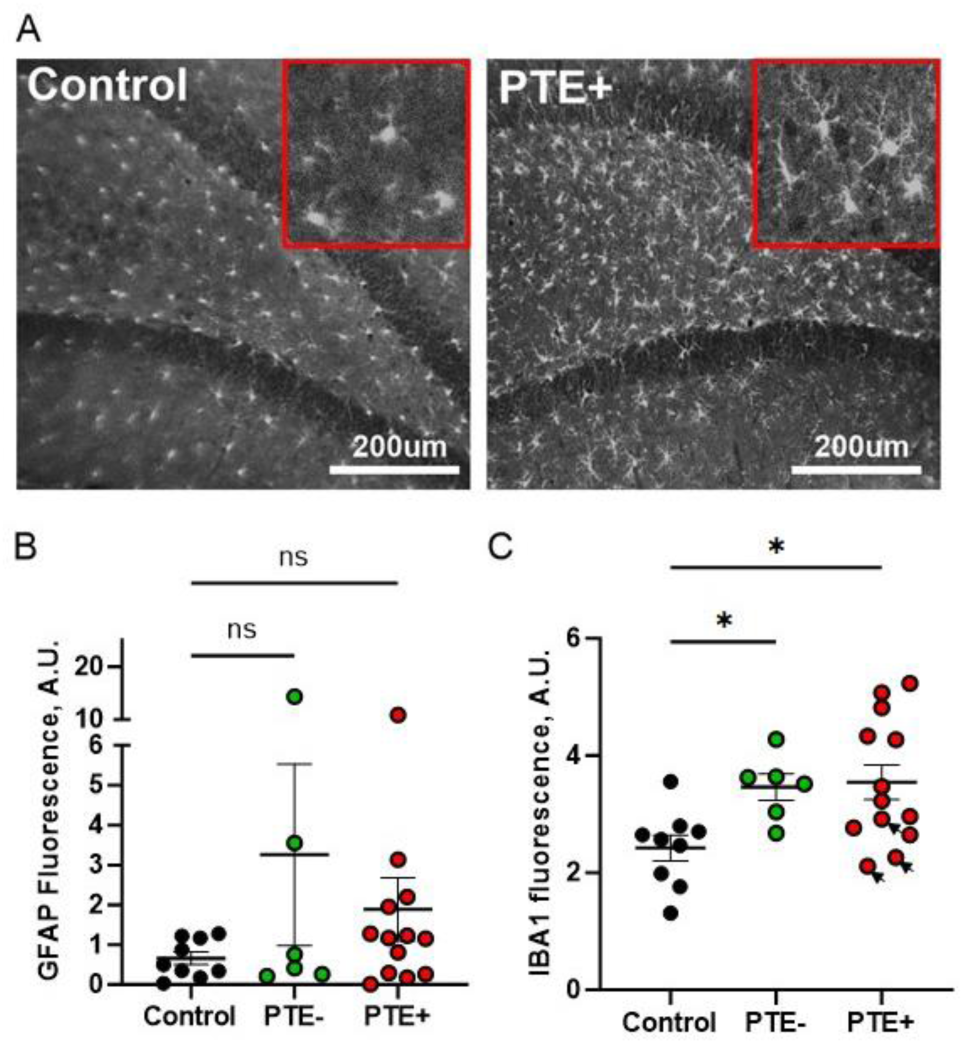
Histological assessment for neuroinflammation. **A)** Representative images showing IBA-1 expression in the dentate gyrus of sham control and PTE+ animals. Insets display magnified views. Scale bar = 200 µm. **B)** No significant differences were found in GFAP expression between groups (Sham control n=9 vs PTE- n=6, P=0.53; Sham control n=9 vs PTE+ n=13, P=0.85; Kruskal-Wallis test followed by Dunn’s post-hoc test). **C)** At 6 months post-injury, elevated IBA-1 expression was observed in PTE+ but not in PTE- animals when compared to sham control animals (Sham control n=9 vs PTE- n=6, P=0.028; Sham control n=9 vs PTE+ n=13, P=0.022; Kruskal-Wallis test followed by Dunn’s post-hoc test). Datapoints from three animals that were epileptic, but no longer showed seizures at time of tissue collection are indicated by arrows.

### PSWEs show diagnostic power for PTE in humans

To test whether the results obtained from rmTBI rats can be translated to brain trauma patients, we analyzed EEG recordings obtained from TBI patients who developed epilepsy after brain trauma. Time in PSWEs (percentage of recording marked with PSWE occurrence) was significantly higher in patients with definitive epileptic EEGs compared to those who were admitted for EEG monitoring to confirm PTE, but their EEG did not reveal epileptiform activity and was within normal range (%64.53 ± %3.15 vs. %17.87 ± %2.89; Fig. 8A). ROC analysis revealed a significant diagnostic power of time in PSWEs for separating definitive epileptic EEGs in our cohort (AUC=0.93; Fig. 8B). Additionally, the elevation of PSWE frequency in patients with confirmed epilepsy demonstrated a generalized distribution across 19 electrodes. Notably, frontal, central and occipital electrodes alike showed a highly significant difference between the two groups. (P<0.0001; Fig. 8C).

**Figure 8.**
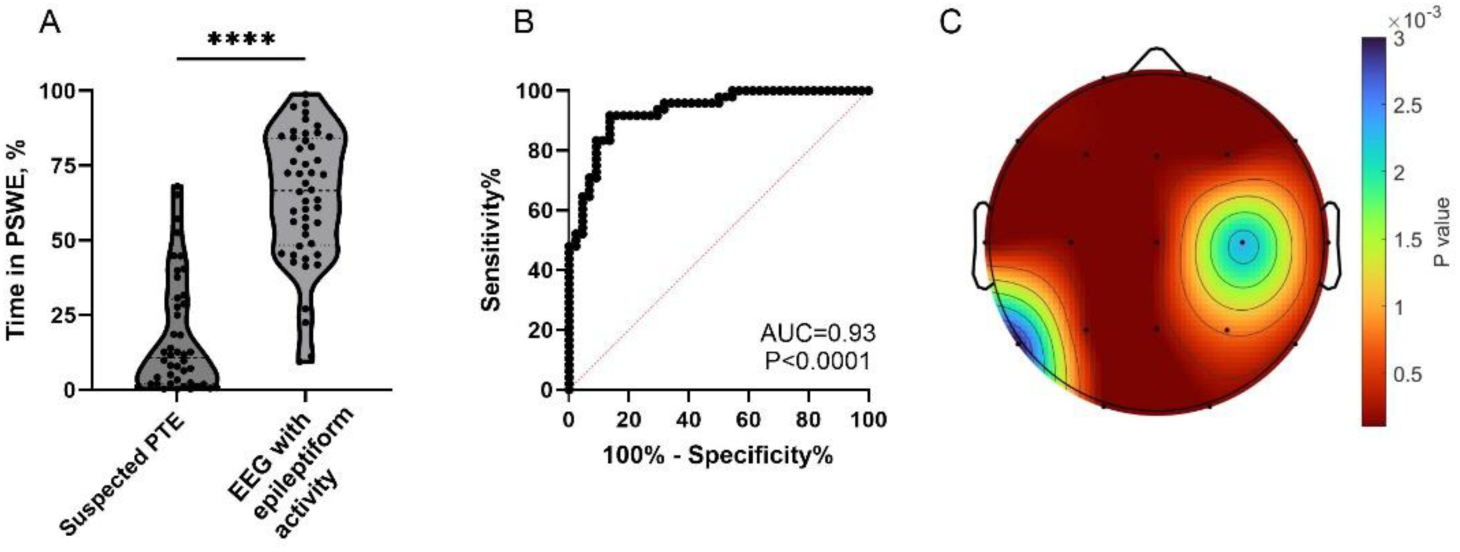
PTE patients with epileptiform EEGs show increased time in PSWEs. **A)** Percentage time in PSWEs was significantly higher in PTE patients with confirmed epileptic EEGs compared to those undergoing EEG monitoring for suspected PTE, where no epileptiform activity was detected, and the EEGs were normal (Suspected PTE n=44 vs EEG with epileptiform activity n=48, P<0.0001; Mann Whitney test). **B)** Time in PSWEs showed significant diagnostic power for separating epileptic EEGs from non-epileptic ones in the ROC analysis (AUC= 0.93, P<0.0001). **C)** Topographical heatmap comparing the mean percentage of time in PSWEs across 19 electrodes between the two groups (Šídák’s multiple comparisons test, 2way ANOVA, P<0.05).

## Discussion

TBI is known to increase the chance of developing a variety of brain pathologies in the weeks to years after head trauma, including neurodegenerative disorders and post-traumatic epilepsy. Intervention might be possible, but knowing which TBI patient is susceptible to developing delayed pathology requires reliable biomarkers. Therefore, we tested acute and early behavioral hallmarks of TBI as a sign for PTE in a rat rmTBI model as well as ECoG and EEG recordings from rats and humans.

The first sign of pathology after head impact was the occurrence of convulsions, which occurred within minutes after injury. These convulsive movements, however, do not necessarily indicate the occurrence of a seizure, but may in fact indicate non-spreading depression of activity or spreading depolarization *(29)*. Both seizures and spreading depolarizations can mediate secondary brain injury, which involves mitochondrial dysfunction *(30)*, neuroinflammation *(26)*, and blood-brain barrier dysfunction *(26, 31)*, which in turn can contribute to delayed pathology and PTE *(32)*. Longer righting latencies following TBI also suggest spreading depolarization *(26)*. These acute convulsive movements, classified as early seizures, were previously linked to disability and PTE in TBI patients *(33)*, which we did not find in our rat rmTBI model. In accordance with previous work in our lab *(26)*, we also found a bimodal distribution in NSS scores in the week of rmTBI.

To test cognitive changes after rmTBI, we measured memory using NOR and anxiety using EPM. Prior studies regularly reported memory decline following TBI in rodents *(34, 35)* and although we did not find significant changes between TBI and controls at 2 months after injury, NOR performance by TBI animals was worse than sham controls at 6 months post rmTBI, in particular by those that also had developed PTE. We also report increased anxiety behavior in TBI rats at 2 months after injury, similar to previous studies in rodents *(34, 36)* and TBI patients *(37–39)*.

Other than behavioral and cognitive changes, we observed the occurrence of SLEs in the obtained ECoG recordings. Interestingly, also control rats displayed frequent SLEs, but those were detected almost exclusively in one electrode. These focal SLEs could be manifestations of spike-wave discharges, which were previously shown to be naturally occurring in outbred, awake rats *(40)*. Alternatively, neural network disturbances during electrode implant could explain these focal SLEs, although postmortem analysis did not reveal structural differences between controls, non-epileptic and epileptic rmTBI animals (Supp. Fig. 1). In contrast, SLEs recorded simultaneously by two or more electrodes, and longer than 20s in duration (IASLEs), showed a more pronounced difference between rmTBI and control animals and most IASLEs also manifested as generalized seizures upon video assessment *(41)*. The average IASLE frequency increases significantly from 2 months to 6 months after injury, which is similar to a previous study in mice *(42)*. Interestingly, the increase in seizure frequency was predominantly driven by IASLEs in the frontal electrodes, which mimics PTE in humans *(43, 44)*. Propagation from frontal to parietal cortex at a velocity of approximately 0.67 mm/s is also in accordance with previous studies *(45)*, although others have reported slightly faster propagation speeds *(46)*. Furthermore, the recorded IASLEs predominantly occurred during the rat’s active phase. It has been well described that seizures can follow a circadian rhythm in epilepsy patients *(47, 48)* and the time of day as well as wakefulness are associated with seizure type and localization *(49)*. In humans, frontal lobe seizures are typically occurring during the night *(49)*, highlighting that the circadian rhythm of rats is hardly translatable to humans.

Upon identifying animals with PTE, we also found high frequency of IASLEs in two control rats (out of 9) at 6 months, which might be due to ageing. More importantly, none of the acute or early behavioral changes could accurately predict PTE. This finding is in line with most previous attempts to use behavior as a predictor of PTE in other rodent models *(50)*, although a previous study reported loss of body weight and slower recovery rate of SNAP scores (combination of several neurological tests) as predictors of PTE *(42)*.

In contrast, PSWE frequency at 2 months after rmTBI was a reliable predictor of developing PTE between 3 and 6 months. PSWEs are characterized as a slowing of brain activity with a median power frequency (MPF) lower than 5 Hz for at least 5 seconds in duration. Increased numbers of PSWEs were previously reported in animal models of epilepsy and in epilepsy patients *(51)*. Increased numbers of PSWEs were also found in aged mice (18–22-week-old) compared to young mice (12-week-old). PSWEs are not associated with all forms of brain pathology or neurodegeneration though. For example, PSWEs have been shown to be uncommon in Parkinson’s disease *(52)*. In our study, PSWEs did correlate with NOR, but not with EPM performance.

Our findings show a significant increase in IBA1 expression in the dentate gyrus of TBI animals, indicating microglial activation and neuroinflammation. Elevated neuroinflammation plays a crucial role in increasing neuronal excitability following TBIs *(54)* and microglial activation is linked to epileptogenesis *(53)*. Here, we report increased IBA1 expression in both PTE- and PTE+ animals. However, three animals that had suffered PTE at earlier timepoints (and thus were identified as PTE+), had seen a significant decline in seizure frequency at the time of tissue collection and could be characterized as seizure-free. Interestingly, these three animals also showed IBA1 expression similar to controls, which suggests that seizure frequency might subside when the inflammatory state is reversed. The localized increase in IBA1 in the hippocampus suggests that neuroinflammation is connected to the seizure-onset zone *(55)* and associated cognitive decline.

PSWEs, which correlate with BBB dysfunction and neuroinflammation, have been shown to be reliable biomarkers for epileptogenesis *(24)*. In the EEG recordings from humans with confirmed epileptiform activity, we found an elevated percentage time in PSWEs, consistent with our animal model. It is worth noting that PTE patients with a high number of PSWEs also show slowing of brain activity consistent with EEG slowing seen in TBI patients with cognitive impairment *(56)*. However, to our knowledge, cortical slowing has not yet been established as a diagnostic nor predictive marker for PTE. Furthermore, a previous study has shown that PSWEs in patients who have experienced a seizure can predict the likelihood of future seizures and thus development of epilepsy *(25)*. Therefore, we propose PSWEs as a promising diagnostic feature, alongside EEG slowing, to identify patients at risk of PTE following brain trauma.

In conclusion, our findings demonstrate that PSWEs may serve as a cost effective, non-invasive reliable predictor of PTE development in both an animal model of rmTBI and human patients, offering a potential biomarker for early intervention.

### Limitations

The present study has several limitations that warrant acknowledgment. First, the use of a single animal model may not fully recapitulate the complexity of TBIs and PTE in humans. Factors such as sub-concussive TBIs, individual variability, pre-existing medical conditions, and inherent anatomical and physiological differences between species are major considerations before generalizing animal data to human populations. Next, despite our efforts to minimize brain damage, the surgical procedures employed in this study still represent an external trauma to the brain, which may explain the high number of SLEs detected in controls (Fig. 2B). Currently however, there are no available non-invasive methods to record brain activity in animal models. Regarding human data, our dataset predominantly included single recordings from patients with a history of PTE. Furthermore, most patients had experienced only a single TBI, or there were no clear indications of repeated brain trauma. Follow-up EEG recordings remain crucial to verify whether PTE patients with high PSWE continue to develop seizures or begin experiencing seizures if they previously did not. Moreover, a more comprehensive data collection from patients with repetitive TBIs would strengthen the conclusions drawn from this study.

## Material & Methods

### Animals

All procedures were approved by the Institutional Animal Care and Use Committee and were performed in accordance with Public Health Service Policy on Humane Care and Use of Laboratory Animals (Protocol no. 20-013). Eight-week-old, wild-type male Sprague-Dawley rats were double-housed in standard cages and exposed to a reversed 12:12 light-dark cycle. All experimentation (Fig. 1A) was performed during the dark (active) phase, except ECoG recordings obtained continuously.

### Weight drop TBI model

rmTBI was induced using a weight-drop model as described (see supplements for the TBI apparatus) *(26)*. In brief, 9- to 12-week-old, male Sprague Dawley rats (n=80) were sedated using an induction chamber (3% isoflurane, 2 L/min O_2_), until the toe-pinch reflex was absent. Rats were then placed in the prone position on a sheet of aluminum foil taped to the top of a plastic box (30x30x20 cm in depth). A metal bolt (1 cm diameter x 10 cm length) was placed on the rat’s head anterior to the lambda suture line (but posterior to bregma) in the midline via alignment with the animal’s ears as an anatomical reference. TBI was induced using a weight (500g) travelling vertically for 0.85m along a metal guide rail, impacting the metal bolt, transmitting the energy onto the rat’s head. Following the impact, animals fell through the foil onto a foam pad placed at the bottom of the box, causing a rotation of the head and neck. Animals were immediately transported to a recovery cage and videotaped during recovery. Weight and distance were set as previously reported *(57)* to result, after a single hit, in a “mild” brain injury which fit the following criteria: (1) neurological deficits following a single injury were transient and reversible within 24 hours; (2) imaging and post-mortem analysis showed no gross brain pathology or intracranial bleeding; and (3) no mortality was observed. The Animal’s acute response to injury was assessed by monitoring involuntary motor convulsive movements, time for regain of righting reflex and neurological severity scores at 24 hours post each hit (see below). Sham controls were anesthetized but did not receive injury. Rats received mild TBI or isoflurane once per day for 4 days.

### Assessment of neurological severity scores (NSS)

Twenty-four hours after each mild TBI, animals were weighed and NSS testing was performed, as described *(26)*, via a composite of open field, beam walk and inverted mesh test scores (four points each). In the open field test, the number of corners explored by the animal within 45 seconds is counted, with a maximum score of 4. For the beam walk test, the animal is placed on a beam with a dark box located on the other size of the beam. The animal is trained for 3 consecutive days to cross the beam successfully. The beam is subdivided in 4 parts of equal length and the distance covered by the animal within 60 seconds is scored, ranging from 0 for no movement, to 4 points for complete crossing. For the inverted mesh test, the animal is placed on top of a mesh and it is immediately turned over. The animal may hang on the mesh grid and the score is determined based on the time until the animal releases from the mesh (0 for immediate release; 1 for <2 seconds; 2 for <4 seconds; 3 for >4 seconds; 4 when the animal climbs back on top of the mesh). NSS scores were determined via video analysis by two blind observers.

### Cognitive assessment

Three weeks and 24 weeks following the rmTBIs we administered elevated plus maze (EPM) and novel object recognition (NOR) adapted from previously established protocols *(58, 59)* to evaluate anxiety-like behavior, learning and recognition memory. EPM anxiety index was calculated as: Anxiety index = 1- [(time in open arm/total time in maze) + (no. entries to open arms/total time in maze)] / 2. Anxiety index values range from 0 to 1, with higher indexes suggesting more anxiety-like behaviour *(59)*. NOR discrimination index was calculated as: Discrimination index = [(no. visits to novel obj.) - (no. visit to old obj.)] / (no. visits to novel obj. + no. visits to old obj.)

### Surgical procedure and Video-ECoG recordings

Four weeks following rmTBI or sham procedures, a subset of animals (Sham control n=9, rmTBI n=23) underwent implantation with electrocorticography (ECoG) recording electrodes. Unipolar, epidural electrodes (Rodent PACK Elec_P1_SS_U, EMKA technologies, France) were implanted in the skull and fixed onto the skull using resin (#651006 Caulk Orthodontic Resin, DENSPLY CAULK) *(60)*. These electrodes were implanted in both hemispheres over the frontal lobe and parietal lobe, together with one reference electrode onto the caudal occipital lobe (see Fig. 1A). Subsequent to the electrode implantation and fixation, the skin was closed with surgical sutures (Prolene BV130-5, ETHICON Inc.) and the animal is allowed to recover.

Following a 1-week post-surgery recovery period, the animals were transported to customized cages for wireless telemetry and video monitoring (Telemetry Rodent PACK system, emka Technologies, France). Lightweight untethered telemetry transmitters (Rodent PACK_TR, emka Technologies, France) were fixed to the animals’ headpieces, allowing for wireless ECoG recording from freely moving animals. ECoG signals were continuously recorded for 48 to 72hrs each month up until 6 months after rmTBI, while video recordings were collected simultaneously. At six months post injury, three rmTBI animals were excluded from the study because they lost their recording headcaps.

### Seizure detection and PSWE analysis

An automated seizure detection algorithm was developed using MATLAB (R2023b, MathWorks, MA, USA). In brief, an artificial neural network (ANN) was trained on expert-verified seizures from various epilepsy models. A researcher-friendly interface (NVision; Biomedical and Health Sciences, Ben Gurion University of Negev) was developed to facilitate its use. For each ECoG recording, two observers, blinded to experimental conditions, randomly selected 60-second-long segments without epileptiform activity to establish a normalization baseline for the detection software. The algorithm then detected “seizure-like events (SLEs)” when the ANN output exceeded a threshold of 1 for at least 6 seconds. The normalisation baseline segments were termed as “SLE-free”.

Behavioral manifestations associated with SLEs and SLE-free segments were assessed by two blinded observers. The observed phenotypes were systematically categorized into three distinct classifications: absence-like staring episodes (stare), minor myoclonic jerks (minor), and major convulsive movements (major). This classification scheme was adapted from standardized epilepsy guidelines to ensure consistency and comparability with existing literature *(61)*.

For PSWE analysis, a modified version of previously established MATLAB algorithm was used to detect cortical slowing and PSWEs in animal and human recordings *(24)*. A PSWE was identified when the MPF dropped below 5 Hz and persisted for a minimum duration of 10 seconds. PSWEs were identified when simultaneous occurrences were detected in at least two recording channels.

### Tissue preparation and immuno-histological analysis

At 6 months post rmTBI, animals were deeply anesthetized with sodium pentobarbital (100 mg/kg, intraperitoneal injection) for transcardial perfusion (Sham control n=9, rmTBI n=19). Physiological saline was first perfused to flush out the blood, followed by 4% paraformaldehyde (AC416785000, Fisher Scientific) for tissue fixation as previously described *(26, 62)*. Brains were cryoprotected using sucrose (15-30%) and coronal brain sections of 30 μm thickness were cut using a freezing microtome (Microm HM400). Free floating sections were immunostained using α-GFAP (1:2000 dilution, Sigma #C9205), α-Iba-1 (1:2000 dilution, Wako #019-19741), α-Rabbit Alexa Fluor488 (1:500 dilution, Invitrogen #A21206) and α-mouse Alexa Fluor568. Image acquisition was performed using a Zeiss Axiocam 503 microscope or LSM880 confocal microscope (Leica SPI 3D STED, Germany), and subsequent quantitative analysis was carried out using ImageJ software (National Institutes of Health).

### Retrospective Analysis of the Temple University Hospital EEG Data Corpus

Human EEG data were obtained from the Temple University Hospital EEG Data Corpus. Outpatient EEG recordings were selected for participants with PTE, identified based on explicit mentions of epilepsy development following brain trauma in their clinical reports. In brief, we employed regular expressions (RegEx) to flag reports containing the keywords "traumatic brain injury," "TBI," "concussion," "head injury," "motor vehicle crash (MVC)," or "motor vehicle accident (MVA)." Further filtering for PTE cases was done using RegEx to identify terms i.e. "seizures," "epilepsy," or "post-traumatic epilepsy." A total of 92 patients were identified following the extraction process, and clinician reports were manually verified. Additionally, EEG impressions and conclusions from epileptologists were assessed to classify patients with definitive epileptiform activity during their recording sessions.

### Statistical analysis

Statistical analyses were conducted using GraphPad Prism 9 (GraphPad Software, San Diego, CA, USA). Significance was assessed using various tests: unpaired or paired t-tests for normally distributed data, Mann-Whitney tests for non-parametric data, Chi-square tests for categorical data, one-way ANOVA with Dunn’s post-hoc tests, two-way ANOVA with Sidak’s multiple comparison tests, and linear regression. Significance was defined as p<0.05. Mean values are reported as Mean ± SEM. Receiver operating characteristic (ROC) analysis was employed to evaluate predictive capacities. Detailed statistical tests and p-values are provided in the figure captions.

## Author contributions

**G.H.**: Formal analysis, Visualization, Validation, Writing – original draft, Writing – review & editing; **P.M.**: Investigation, Formal analysis, Visualization, Writing – original draft, Writing – review & editing; **H.I.**: Software, Formal analysis; **E.P.**: Methodology **S.M.**: Investigation, Formal analysis; **L.A.H.**: Investigation; **M.A.**: Formal analysis; **Z.A.**: Formal analysis; **A.F.**: Conceptualization, Writing – review & editing, Supervision, Funding acquisition.

## Acknowledgements

We thank Kathleen Murphy and Jim Kukurin for their technical expertise.

## Supplementary Material

**Supplementary figure 1.**
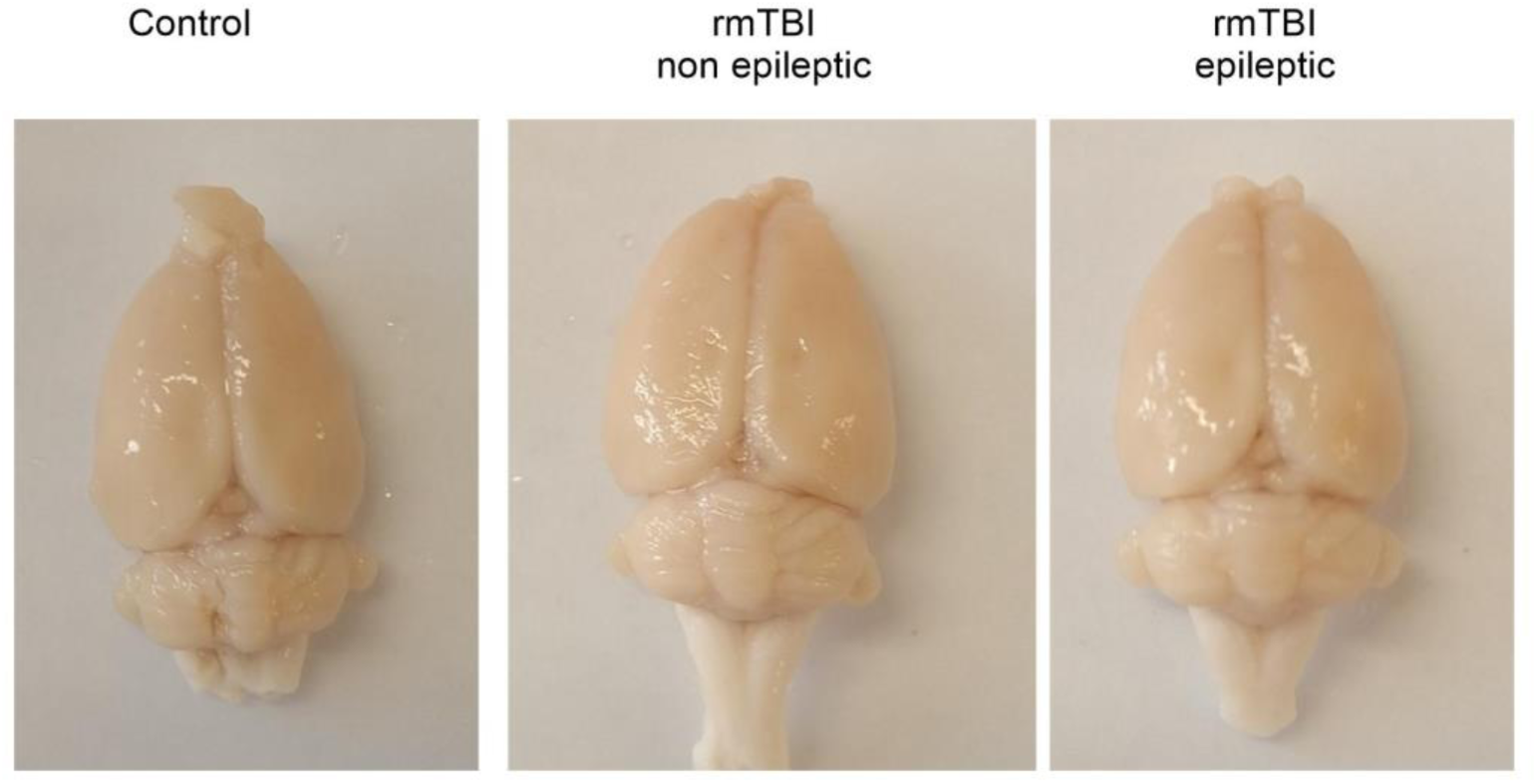
No gross anatomical differences were observed in brains at six months post injury between groups.

